# Capturing the signalling dynamics of the MAPK-AKT-mTOR pathway in a single targeted phosphoproteomics assay

**DOI:** 10.1101/2022.01.17.476555

**Authors:** Donna O. Debets, Juan Manuel Valverde, Maarten Altelaar

## Abstract

The MAPK-AKT-mTOR protein network integrates extra- and intracellular signals to determine cellular fate, regulating pivotal biological processes such as cell growth and metabolism. Due to this crucial role, pathway dysregulation has been implicated in multiple diseases, such as metabolic disorders and cancer. The MAPK-AKT-mTOR pathway consists of dozens of proteins and signal transduction is primarily driven by protein phosphorylation. Here, we present a targeted phosphoproteomics assay to study the phosphorylation dynamics of the MAPK-AKT-mTOR pathway in detail with high sensitivity and in a high throughput manner. By using a multi-protease approach, we increased the pathway coverage with phosphosites that were previously inaccessible. This novel approach yields the most comprehensive method for the detailed study of mTOR signalling to date (covering 150 phosphopeptides on more than 70 phosphoproteins), which can be applied to *in vitro* and *in vivo* systems and has the sensitivity to be compatible with small sample amounts. We demonstrate the feasibility of this assay to monitor the plasticity of MAPK-AKT-mTOR phosphorylation dynamics in response to cellular stimuli with high temporal resolution and amino acid residue specificity. We found highly dynamic phosphorylation events upon treatment with growth factors, revealing the sequential nature of phosphosites in this signalling pathway. Furthermore, starvation of glucose and amino acids showed upregulation of AKT-targets PRAS40^T246^ and FOXO3^T32^, highlighting the role of AKT in cellular response to starvation. These findings illustrate the potential of this assay to obtain new biological insight when monitoring dynamics of functional phosphosites.

**Highlights:**

1. Robust targeted MS assay to study the phosphorylation dynamics of the MAPK-AKT-mTOR network
2. Extended pathway coverage by application of multiple proteases for protein digestion
3. Highly sensitive, high throughput and readily applicable assay for *in vivo* and *in vitro* systems
4. Phosphorylation patterns of MAPK-AKT-mTOR network are highly dynamic and change upon stimulation with growth factors, amino acids and glucose

**Motivation:** The MAPK-AKT-mTOR protein network integrates extra- and intracellular signals to determine cellular fate, regulating pivotal biological processes such as cell growth and metabolism. Due to this crucial role, pathway dysregulation has been implicated in multiple diseases, such as metabolic disorders and cancer. Our understanding of the complex regulation of this intricate signalling network is incomplete and is hampered by the lack of analytical methods to study its phosphorylation dynamics in detail. In this study, we present a targeted phosphoproteomics assay to monitor the phosphorylation events of the MAPK-AKT-mTOR pathway with amino acid residue specificity and in a high throughput manner. We increased the pathway coverage with phosphosites that were previously inaccessible by the use of multiple proteases for protein digestion. This novel approach yields the most comprehensive method for the detailed study of MAPK-AKT-mTOR signalling to date, which can be applied to *in vitro* and *in vivo* human samples and has the sensitivity to be compatible with small amounts of starting material.

**Graphical abstract:** 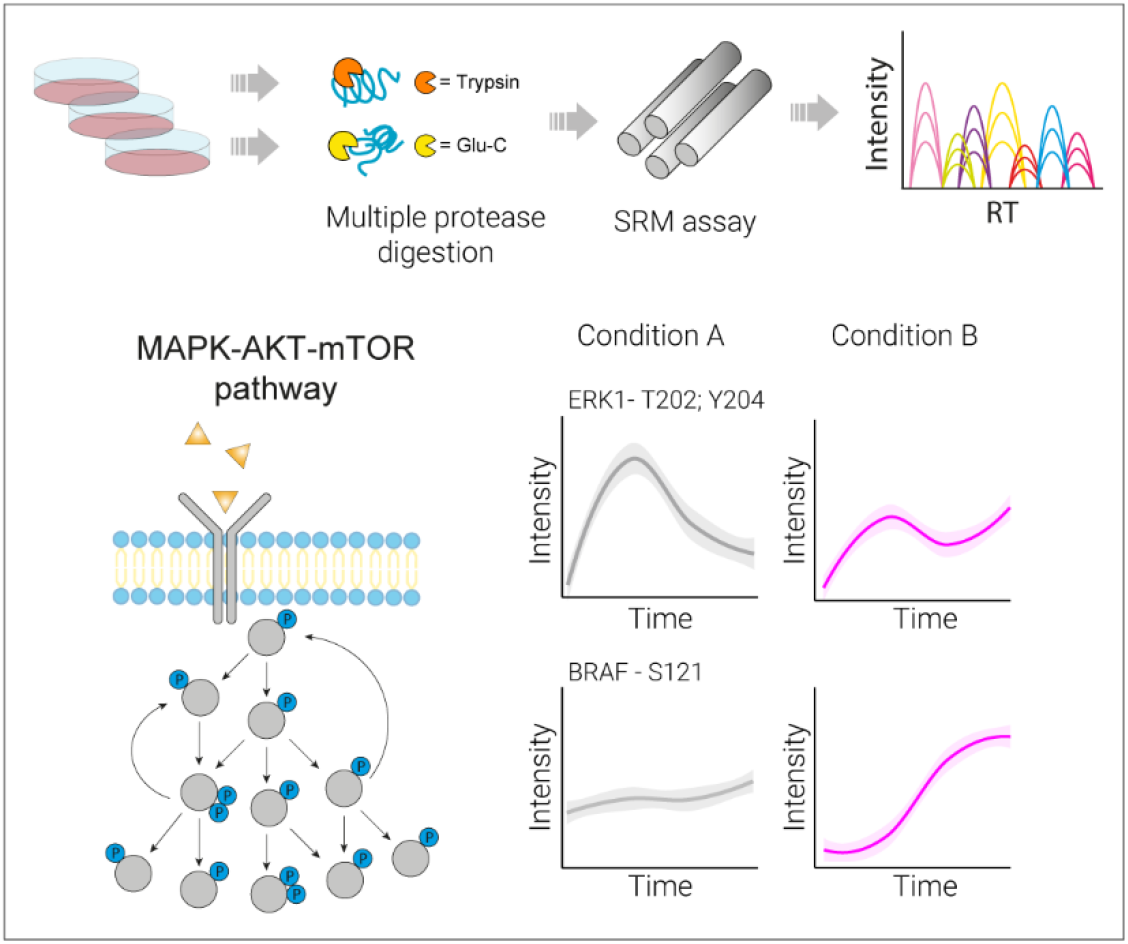

## Introduction

Cells continuously sense extra- and intracellular stimuli to regulate cell growth and metabolism. Under healthy conditions, proliferation is stimulated when conditions are favourable and halted under stress. To achieve this, a wide variety of stimuli, from growth factors and nutrients to DNA damage, are sensed and integrated by signalling pathways to determine cellular fate (Liu and Sabatini, 2020). These pathways are driven by protein phosphorylation, which allows for fast and dynamic transmission of information throughout the cell (Jin and Pawson, 2012) (Hunter, 2012).

In eukaryotes, mitogen-activated protein kinase (MAPK) cascades are key players to relay information from growth factor stimuli at the cell surface to the interior of the cell (Seger and Krebs, 1995). The kinases in these pathways have been heavily studied and are known to be key regulators of processes ranging from embryonic development to cell proliferation and growth (Burotto et al., 2014; Drosten and Barbacid, 2020; Hamilton and Brickman, 2014). In addition to MAPKs, the phosphoinositide 3-kinase (PI3K)-AKT signalling network is another major integrator of growth factor stimuli, like insulin and cytokines (Hoxhaj and Manning, 2020), making it another essential regulator of cell fate. Hence, proteins of both of these pathways are often found mutated in malignancies (Lawrence et al., 2014). Interestingly, these pathways converge into two protein complexes, mTOR complex 1 (mTORC1) and mTOR complex 2 (mTORC2).

Both mTORC1 and mTORC2 contain the mTOR serine/threonine protein kinase, which functions as the catalytic subunit responsible for phosphorylation of other protein kinases within the signalling cascade and effector proteins that directly regulate cell metabolism (Saxton and Sabatini, 2017). Besides the mTOR kinase, each complex has additional subunits that function as either enhancers or inhibitors of its kinase activity. Some of these subunits are directly regulated by MAPK and PI3K-AKT activity (Mendoza et al., 2011). The mTOR network controls processes such as ribosome biogenesis, mRNA translation, cytoskeleton organisation and both catabolic and anabolic programs, like autophagy and lipid biosynthesis, respectively (Liu and Sabatini, 2020; Saxton and Sabatini, 2017). Together, the MAPK-AKT-mTOR signalling network converts extracellular signals (mainly via MAPK and AKT signalling) and intracellular environmental cues (e.g. DNA damage and ATP levels) to determine cellular fate.

The MAPK-AKT-mTOR pathway is composed of dozens of proteins, which transmit signals via phosphorylation of specific amino acid residues on target proteins. Therefore, to elucidate the complex regulation of this intricate signalling network, analytical methods to monitor the dynamics of these phosphosites are required. Protein phosphorylation events can be studied using antibody-based techniques, such as western blotting. These are generally very sensitive, yet their use is restricted by the availability of adequate phosphosite-specific antibodies. Furthermore, western blots are merely semi-quantitative and are difficult to multiplex. These limitations can be overcome by the application of targeted mass spectrometry (MS), namely selected reaction monitoring (SRM) (Adachi et al., 2016). By selectively measuring peptides and peptide fragments of interest, this technique can monitor phosphosites with amino acid residue specificity (de Graaf et al., 2015). Furthermore, combined with highly selective phosphopeptide enrichment, SRM provides high sensitivity. Quantitative reproducibility is ensured using heavy labelled standards. Targeted MS methods are easily multiplexed and allow for hundreds of phosphopeptides to be measured in a single MS run.

Here, we developed a targeted MS assay to study the phosphorylation dynamics of the MAPK-AKT-mTOR pathway in a high throughput manner, selecting phosphorylation sites with annotated biological function. We make use of multiple proteases for protein digestion, which increases the pathway coverage with phosphosites that were previously inaccessible. This novel approach yields the most comprehensive method for the detailed study of MAPK-AKT-mTOR signalling to date, which can be applied to *in vitro* and *in vivo* human samples and has the sensitivity to be compatible with small amounts of starting material. In a proof-of-principle experiment, we demonstrate that we can monitor the great plasticity of MAPK-AKT-mTOR phosphorylation dynamics with high temporal resolution and amino acid residue specificity. We show unique phosphorylation patterns for different stimuli and find sequential phosphorylation events that correlate with their known functionality. Overall, we think this assay could be a great resource for scientists who are interested in obtaining mechanistic insight in how phosphorylation via the MAPK-AKT-mTOR network controls cell growth and metabolism.

## Results

### Selection of phosphopeptides for assay development

We first performed a literature search to identify the main phosphoproteins in the MAPK-AKT-mTOR pathway and subsequently selected the principal phosphosites on these proteins, focussing on sites with known functionality. Next, we aimed to choose the most suitable phosphopeptides for LC-MS/MS analysis for each phosphosite of interest (Figure 1A, top). We therefore generated an extensive phosphopeptide library using BJ-EHT fibroblast cells transduced with an inducible RAS mutation. Induction of RAS activity in this cell line results in hyperphosphorylation of the MAPK pathway and hence enhanced our ability to detect phosphosites of interest. To extend the coverage of the phosphoproteome beyond sites accessible by trypsin digestion, proteins were digested in parallel by four proteases: Trypsin/LysC, AspN, GluC or Chymotrypsin. The generated peptides were fractionated by high pH fractionation prior to phosphopeptide enrichment and LC-MS/MS analysis (Figure 1A, bottom).

**Figure 1.**
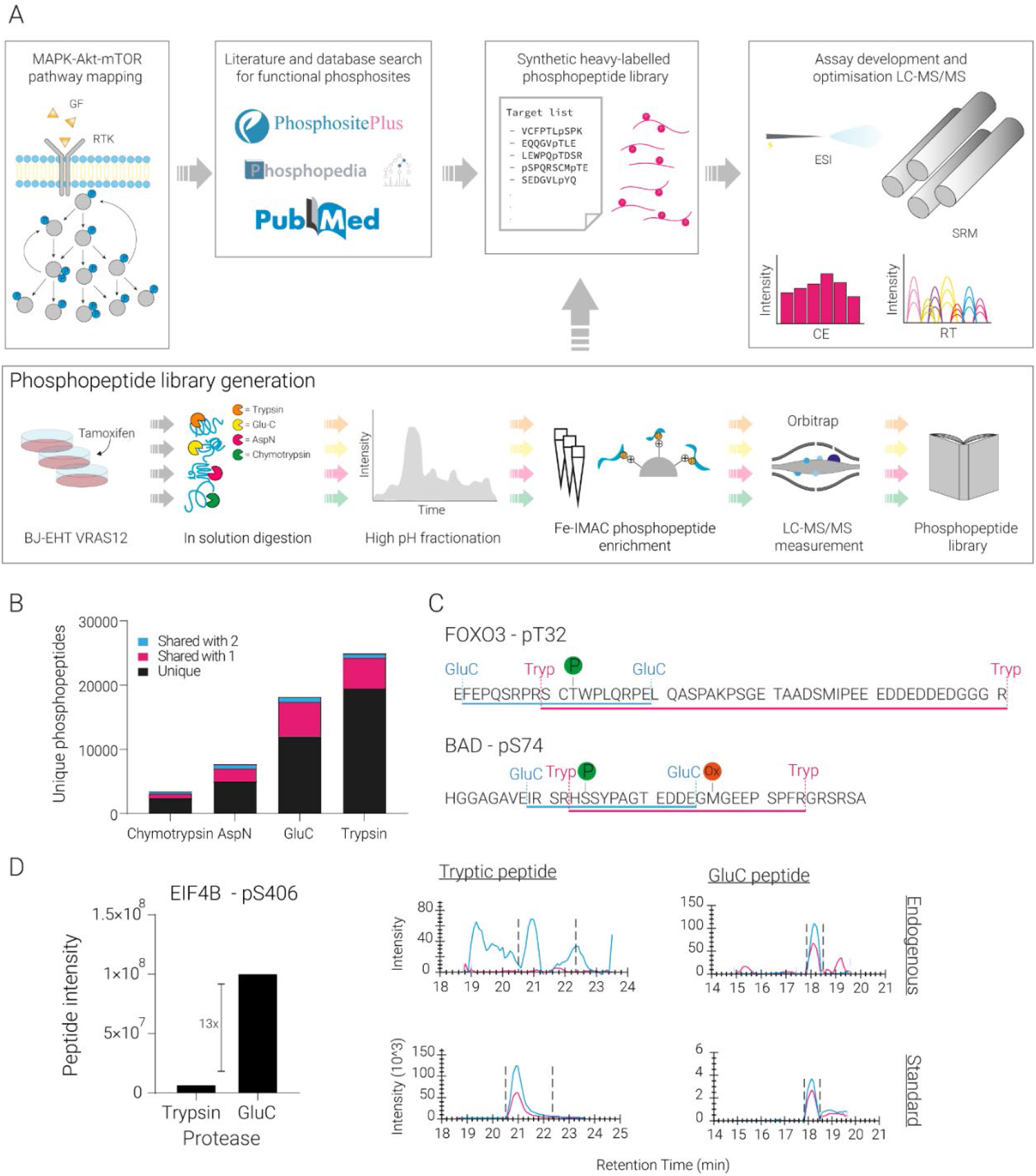
Development of targeted phosphoproteomics assay and multi-protease approach. (A) Top: Mapping of proteins in the MAPK-AKT-mTOR network was done by comprehensive literature mining. Functional phosphosites were selected and information on the corresponding phosphopeptides was obtained from our phosphopeptide library (bottom panel) combined with Phosphopedia. The selected sequences were synthesised, incorporating heavy labelled amino acids, and used as internal standards and for assay development. Collision energy (CE) was optimised for each transition individually to increase sensitivity. Retention time scheduling was performed to measure hundreds of peptides in one MS run. (A) Bottom: An in-house phosphopeptide library was created using cells with inducible RAS activity, which were subjected to lysis and protein digestion with multiple enzymes in parallel (Trypsin/LysC, GluC, AspN and Chymotrypsin). High pH fractionation was used to maximize phosphopeptide detection, followed by Fe-IMAC enrichment and MS acquisition. The generated phosphopeptide library was mined for phosphopeptides of interest within the MAPK-AKT-mTOR pathway. (B) Number of phosphopeptides per protease. (C) Example of a phosphosite that was less favourable by trypsin digestion (top): tryptic peptide for FOXO3^T32^ was too long for efficient MS detection. Example of a phosphosite with favourable peptide properties using GluC for digestion (bottom): The GluC peptide for BAD^S74^ lacks the methionine, which is prone to oxidation during sample preparation. (D) GluC digestion improved detection of endogenous EIF4B^S406^. Higher intensity for endogenous EIF4B^S406^ by GluC compared to trypin digestion was found in our peptide library (left). This resulted in higher sensitivity in the SRM assay (right).

In total, our phosphopeptide library consisted of 46,080 unique phosphosites on 9065 phosphoproteins (Supplementary Figure 1). Importantly, each proteolytic enzyme generated peptides rendering unique phosphosites and phosphoproteins that were not identified by any of the other proteases. Protein digestion with Trypsin/LysC generated the highest number of (unique) phosphosites, followed by GluC, AspN and Chymotrypsin (Figure 1B).

To generate our targeted phosphoproteomics assay, we extracted the biologically relevant phosphosites from our phosphopeptide library. Since merely three phosphosites of interest were uniquely identified with AspN and Chymotrypsin, these proteases were excluded for further use. Proteolytic digestion with Trypsin/LysC and GluC contrarily generated unique phosphopeptides covering many more phosphosites within the pathway. For example, FOXO3^T32^ was accessible with GluC digestion, whereas tryptic digestion did not generate a peptide suitable for LC-MS/MS analysis (Figure 1C). Additionally, the use of multiple proteases allowed us to select peptides with advantageous physiochemical properties. Phosphopeptides containing a methionine were disfavoured since methionine oxidation can occur *in vitro*, complicating method development and potentially reducing sensitivity since the signal is split over two peaks (Figure 1C). Lastly, we took advantage of the differences in detectability of specific phosphopeptides generated by the two proteases. Endogenous EIF4B^S406^ was detected at a much higher intensity (13x) after GluC digestion compared to Trypsin/LysC. This quantitative benefit resulted in increased sensitivity of our targeted phosphoproteomics assay (Figure 1D). Overall, the application of multiple proteases for protein digestion increased the coverage of biologically relevant phosphosites and allowed for the selection of peptides with suitable chemical and quantitative properties.

### Assay development

In total, we included 150 phosphopeptides covering 72 phosphoproteins in the MAPK-AKT-mTOR pathway, covering many of the biologically relevant phosphosites within this signalling network (Figure 2A). All selected phosphopeptides were synthesised incorporating a heavy isotopic label and were used as reference standard. We observed good sensitivity and a large linear range of quantification as demonstrated by dilution series of our synthetic peptides (Supplementary Figure 2A). The quantitative reproducibility was further determined by replicate measurements, which showed good coefficient of variation (Supplementary Figure 2B). To measure all phosphopeptides in a single LC-MS/MS run, we optimised retention time scheduling using a dimensionless value that reflects chromatographic retention relative to a mix of commercial standards (iRTs) (Escher et al., 2012). Therefore, robust LC separation was critical (Supplementary Figure 2C).

**Figure 2.**
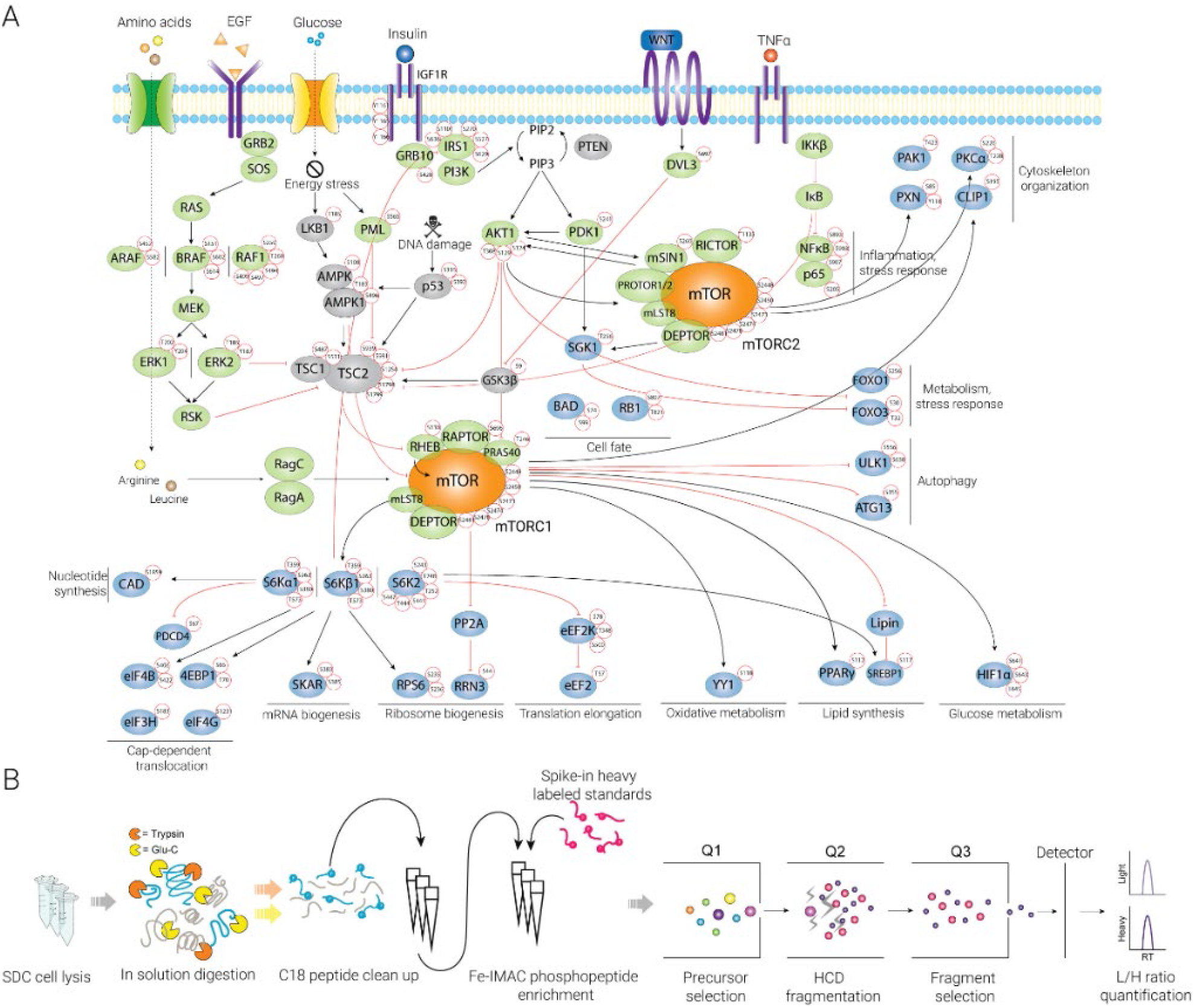
Targeted phosphoproteomics assay extensively covers the biologically relevant phosphosites in the MAPK-AKT-mTOR pathway. (A) Comprehensive view of the protein network, in red circles the phosphosites included in our assay. Green nodes represent activators of mTOR kinase activity, grey nodes represent inhibitors, blue nodes are effector proteins. Black arrows suggest activation, red arrows indicate inhibition. (B) Targeted assay workflow.

In brief: for our targeted phosphoproteomics assay, first cells were lysed and proteins were digested in parallel with either GluC or Trypsin/LysC. GluC and Tryptic peptides were pooled prior to automated sample clean-up. Heavy standards were added prior to phosphopeptide enrichment, and the enriched samples were measured on a nanoLC coupled to a triple-quadrupole MS (Figure 2B).

### MAPK-AKT-mTOR phosphorylation dynamics

To assess the performance of our targeted phosphoproteomics assay, we analysed the MAPK-AKT-mTOR phosphorylation dynamics of MCF10A cells stimulated with EGF and insulin. Cells were starved for 24 hours before stimulation with EGF and insulin, and samples were harvested after 24h starvation and 1 min, 30 min and 60 min after stimulation (Figure 3A). We also assessed MAPK-AKT-mTOR phosphorylation status under normal growth conditions (24 hours in fully supplemented media). Cell growth was halted under starvation, but rescued upon addition of EGF and insulin (Supplementary Figure 3). In total, we quantified 55 phosphorylation events across the different conditions. These phosphorylation events displayed distinct trends, grouping phosphopeptides into five different clusters (Figure 3B). The first three clusters represented phosphosites that were downregulated during normal growth conditions and starvation but were clearly responsive to addition of EGF and insulin after starvation. Within these three clusters, we could discern phosphorylation events that were instantly upregulated after growth factor addition (within a minute) and swiftly downregulated (cluster 1, ‘early response’), behaving almost in a switch-like fashion. Furthermore, we identified a group of phosphosites that were quickly upregulated and remained upregulated over a longer period (cluster 2, ‘sustained response’). Lastly, cluster 3 represented phosphorylation events that were upregulated upon growth factor addition at a later stage (30-60 min) and remained upregulated during the time course of the experiment (‘delayed response’). These data highlight the feasibility of our assay to detect dynamic phosphorylation events upon stimulus.

**Figure 3.**
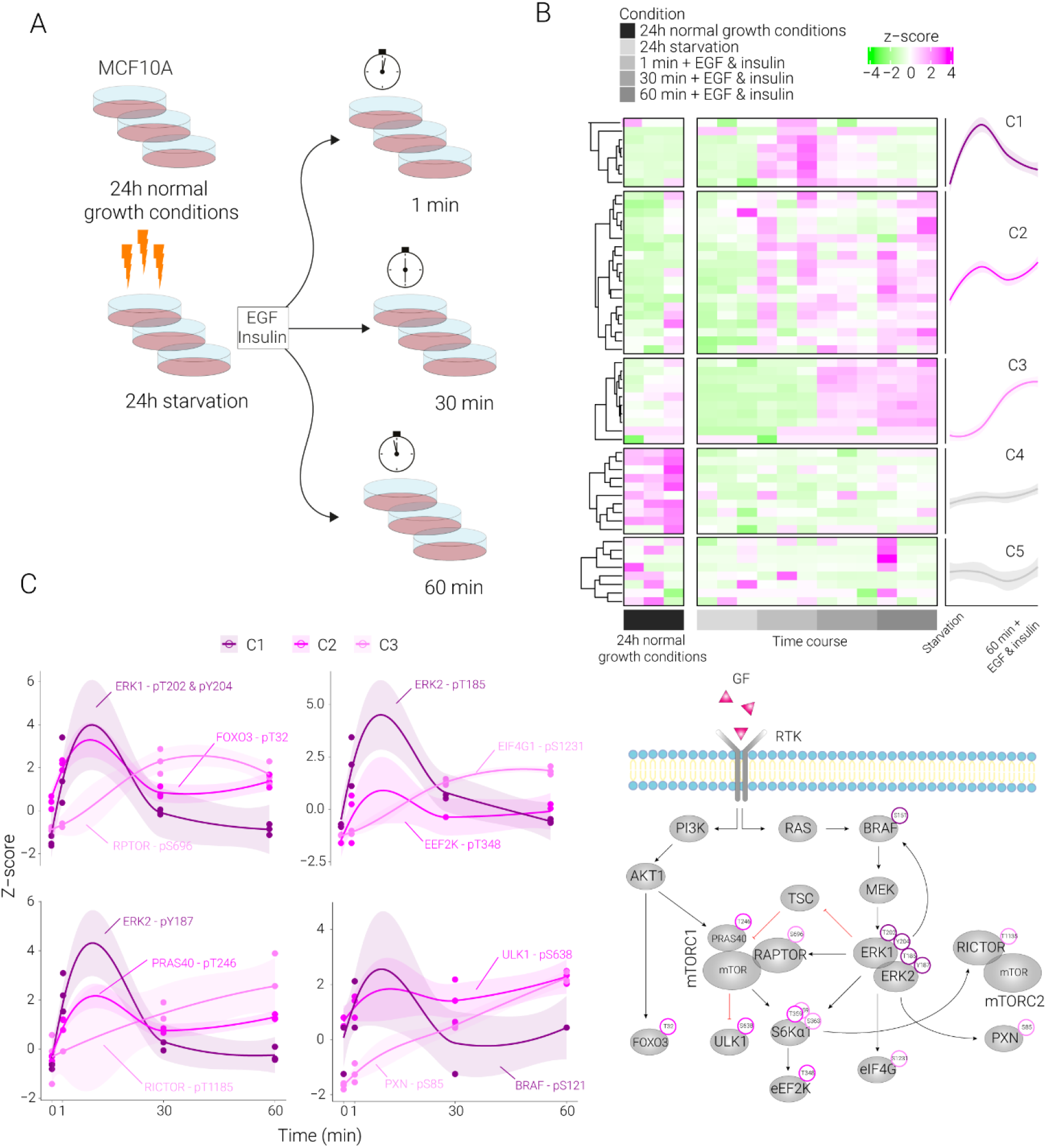
Probing MAPK-AKT-mTOR activity after growth factor stimuli. (A) Experimental setup: MCF10A cells were grown under normal conditions for 24hours, followed by starvation in additive-free medium for 24 hours, after which cells were stimulated with EGF and insulin. Cells were harvested at indicated time points. (B) Unsupervised clustering of quantified phosphopeptides highlights five clusters with distinct phosphorylation patterns. (C) Profile plots of the dynamics of regulated phosphosites (left) correlating with suggested cascade of signalling events (right). Colour represents the clusters in Figure 3B.

In contrast to cluster 1, 2 and 3, the phosphorylation events in clusters 4 and 5 were not responsive to addition of EGF and insulin. Interestingly, the phosphosites in cluster 4 were upregulated during normal growth conditions (full media), decreased upon starvation and did not recover after replenishing with EGF and insulin (Figure 3B). Multiple phosphosites in this cluster were directly involved in protein translation, namely EIF4B^S406^, EIF3H^S183^ and 4EBP1^S65^. EIF4B and EIF3H are crucial components of the translation initiation complex, and their functionality is phosphorylation-driven; increased phosphorylation is positively correlated with induced translation (Holz et al., 2005; van Gorp et al., 2009). Furthermore, the hypophosphorylated form of 4EBP1 represses translation initiation by binding to initiation factor EIF4E and hence inhibiting its assembly into the translation initiation complex (Gingras et al., 1999). Downregulation of these three phosphosites results in reduced translation, which is anticipated upon starvation. Interestingly, phosphorylation translation initiation factor eIF4G1^S1231^ was found upregulated in cluster 3 (‘late response’). This phosphosite is known to be regulated by ERK and correlates with formation of the initiation complex (Dobrikov et al., 2013). Overall, our data indicate that addition of EGF and insulin after prolonged starvation did not restore protein translation within this relatively short timeframe (60 min).

Next, we set out to establish whether we could identify any kinase-substrate pairs from the dynamic phosphorylation events in cluster 1 (‘early response’), cluster 2 (‘sustained response’) and cluster 3 (‘delayed response’). Indeed, in cluster 1 we identified phosphorylation of mitogen-activated protein kinases, namely BRAF^S121^, ERK1^T202/Y204^ and ERK2^T185/Y187^, these kinases are known to be readily activated upon growth factor stimulation. Phosphorylation of BRAF^S121^ by ERK activates a negative feedback loop inhibiting the interaction between RAS and BRAF, resulting in reduced phosphorylation of ERK (Ritt et al., 2010). This could explain the transient phosphorylation of ERK we observed (Figure 3C).

Increased ERK activity directly following stimulus (cluster 1), resulted in phosphorylation of ERK substrates at later time points (cluster 3) (Figure 3C). Amongst these was RPTOR^S696^, which upon upregulation activates mTORC1, resulting in further downstream phosphorylation of kinase S6KA1^T359/S363^ (Figure 3C). S6KA1 has been reported to phosphorylate RICTOR^T1135^, bridging the mTORC1 pathway with mTORC2 signalling. These phosphorylation dynamics are in concordance with the cascade of phosphorylation events previously reported in literature (Figure 3C).

In cluster 2, we identified phosphorylation events indicative of AKT activation upon insulin stimulation: upregulation of FOXO3^T32^ and PRAS40^T246^ (Figure 3C) (Kovacina et al., 2003; Vander Haar et al., 2007; Zheng et al., 2000). FOXO3 triggers apoptosis in the absence of survival factors, whereas upon phosphorylation by AKT, FOXO3 is retained in the cytosol, restricting its pro-apoptotic function (Tzivion et al., 2011). PRAS40 on the other hand is a negative regulator of mTORC1. Upon phosphorylation, PRAS40 dissociates from mTORC1, activating mTOR signalling (Wang et al., 2007). The increased and lasting upregulation of FOXO3 and PRAS40 phosphorylation upon growth factor stimulation found in this experiment shows activation of the PI3K-AKT pathway resulting in reduced pro-apoptotic signals and increased mTORC1 activity. This heightened mTOR activity can subsequently lead to phosphorylation of its downstream substrate, the kinase ULK1^S638^ (Figure 3C). ULK1 activation upon starvation initiates an autophagy response, whereas phosphorylation of ULK1 by mTOR results in inhibition of autophagy when nutrients are replenished (Shang et al., 2011). Our data suggests that activation of mTORC1 upon EGF and insulin stimulation also results in ULK^S638^ phosphorylation. Together, these results highlight the feasibility of our assay to disentangle patterns of sequential phosphorylation within the MAPK-AKT-mTOR pathway.

After highlighting phosphorylation dynamics caused by stimulus with growth factors, we set out to assess the effect of other nutrients known to modulate MAPK-AKT-mTOR activity. For this, we starved cells of either glucose or amino acids, using nutrient free media. Subsequently, we reincorporated each nutrient and collected samples at different time points. Since these experiments involved complete exchanges of media, we performed a control experiment changing media with the same composition and collected samples at the same time points (Figure 4A). Overall, we quantified 39 and 49 phosphopeptides upon stimulation with amino acids and glucose, respectively. Stimulus with these nutrients did not reveal a strong dynamic phosphorylation response, in clear contrast with growth factor treatment observed before (Figure 3B). Interestingly, glucose and amino acid deprivation caused increased phosphorylation of PRAS40^T246^ and FOXO3^T32^, which were both downregulated upon reincorporation of each nutrient. As mentioned previously, both phosphosites were clearly upregulated after treatment with growth factors (Figure 4B). If these two sites are used as proxy of AKT activity, our results suggest that nutrient starvation causes upregulation of PI3K-AKT pathway. Others have previously suggested that upon depletion of glucose and amino acids cells engage on a survival program by increasing the activity of the MAPK-PI3K-AKT pathways, in order to ensure mTOR activation and promote cell survival (Figure 4C) (Pathria and Ronai, 2021). Overall, these results show that our assay can be used to discriminate differences in MAPK-AKT-mTOR signalling upon treatment with different stimuli.

**Figure 4.**
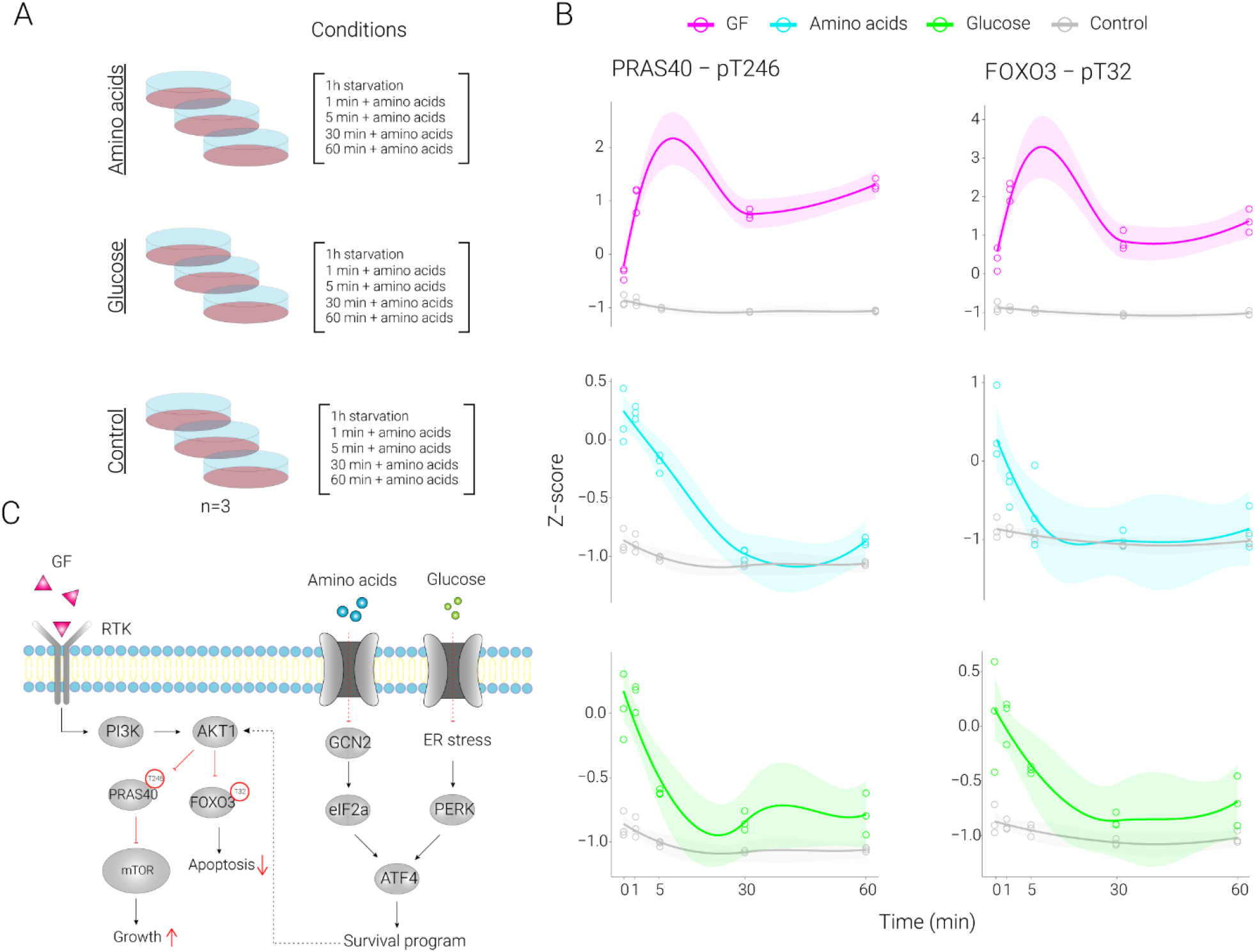
Studying the phosphorylation dynamics of MAPK-AKT-mTOR after stimulus with glucose and Amino Acids following starvation. (A) Experimental setup: MCF10A cells were starved in glucose-free or amino acid-free media for 1 hour, followed by stimulus with glucose or amino acids respectively. Cells were harvested after 1 hour of starvation and at different time points following stimulus. (B) Response of PRAS40^T246^ and FOXO3^T32^ shows differential response after starvation and stimulus with glucose and amino acids, when compared to growth factor (GF) treatment. Time point 0 represents starvation. (C) PRAS40^T246^ and FOXO3^T32^ illustrate AKT activation due to a cellular survival program induced by glucose or amino acid deprivation (Pathria and Ronai, 2021).

## Conclusion & Discussion

In this study, we developed a targeted MS assay to study functional phosphorylation dynamics of the MAPK-AKT-mTOR pathway in detail, with high reproducibility and sensitivity. Even though trypsin is the most commonly used digestive enzyme in proteomics studies due to its favourable characteristics (Vandermarliere et al., 2013), our study highlights the advantage of the use of multiple proteases to extensively cover the important phosphosites within the pathway, which here can be combined in a single MS analysis. The benefit of this approach is especially high for the study of posttranslational modifications since this requires the measurement of specific peptide sequences. This was evident from our phosphopeptide library which included unique phosphosites and phosphoproteins for each protease. Overall, our assay highlights the potential of custom-made targeted proteomics assays to measure specific phosphosites of interest.

In our experiments, we demonstrated the ability to monitor the plasticity of the phosphorylation dynamics in the MAPK-AKT-mTOR network with high temporal resolution and amino acid residue specificity, under diverse external stimuli. We found that most proteins involved in translation initiation were not phosphorylated upon EGF and insulin stimulus, contrary to previous studies (Holz et al., 2005; van Gorp et al., 2009). Only eIF4G1^S1231^ was found upregulated in cluster 3 (‘delayed response’). This suggests that translation initiation factors are not phosphorylated at the same rate. The lack of increased phosphorylation of other initiation factors might be due to the relatively short time course of our experiment or could indicate that replenishing with EGF and insulin (in absence of serum) is inadequate to fully rescue translation initiation after prolonged starvation. Further experiments are needed to fully elucidate the phosphorylation dynamics of translation initiation factors after starvation.

We found dynamic ULK1 phosphorylation upon stimulation with EGF and insulin after starvation. Increased mTOR activity has been shown to phosphorylate ULK^S638^, inhibiting autophagy upon replenishing with full media (Shang et al., 2011). Our data shows that stimulation with EGF and insulin (and the resulting mTOR activation) is sufficient to induce ULK1^S638^ phosphorylation. Additionally, we found upregulation of ULK1^S556^ upon growth factor stimulus. This phosphosite is responsive to AMPK activity upon starvation and increased ULK1^S556^ has been suggested to induce autophagy (Jang et al., 2017; Lin et al., 2021). Therefore, increased ULK^S556^ upon EGF and insulin stimulation is surprising. This discrepancy highlights that the ULK1 phosphorylation dynamics are not fully elucidated.

Next, by using PRAS40^T246^ and FOXO3^T32^ as a proxy of AKT activity, we found evidence in our data of AKT activation upon amino acid and glucose starvation. This is in line with previous research which has proposed increased AKT activity as a survival mechanism upon depletion of these nutrients (Gwinn et al., 2018; Pathria and Ronai, 2021; Shin et al., 2015). We showed that AKT activation of mTORC1 via PRAS40^T246^ and transcriptional regulation via FOXO3^T32^ are potential effectors of this survival program. Interestingly, our results did not show upregulation of the MAPK pathway upon nutrient starvation, which has also been proposed as a survival mechanism (Pathria et al., 2019; Thiaville et al., 2008). This indicates that PI3K-AKT has a more prominent role than MAPK in this type of survival response. This is of interest since both amino acid and glucose starvation have been proposed as therapeutic strategies against certain malignancies (Ganapathy-Kanniappan and Geschwind, 2013; Kang, 2020). Overall, these findings suggest that PRAS40 and FOXO3 could be potential therapeutic targets, which could be exploited by combining nutrient starvation with AKT inhibitors.

In conclusion, our experiments described here highlight the potential of our targeted phosphoproteomics assay to monitor the highly dynamic phosphorylation events of the MAPK-AKT-mTOR pathway. Our method extensively covers the important phosphosites in the network (including 150 phosphopeptides on more than 70 phosphoproteins), is sensitive, high throughput and can be applied to *in vivo* and *in vitro* human samples. Therefore, we believe that this assay is a great resource for scientists to study the integration of environmental cues by the MAPK-AKT-mTOR pathway to regulate cellular fate in health and disease.

## Acknowledgements

We would like to thank Pieter C. van Breugel for providing us with the BJ-EHT fibroblast cells and Mirjam J. Damen for help with the high pH fractionation. This work has been supported by EPIC-XS, project number 823839, funded by the Horizon 2020 programme of the European Union and the NWO funded Netherlands Proteomics Centre through the National Road Map for Large-scale Infrastructures program X-Omics, Project 184.034.019. JMV is supported by scholarships from the Ministry of Science and Technology of Costa Rica (MICITT) and the University of Costa Rica (UCR).

## Methods

### Generation of phosphopeptide library

BJ-EHT vRAS12 cells were cultured in DMEM and treated with 100 nM tamoxifen for 1hr, or 24hr to induce RAS activity. Cells were washed with ice cold PBS, detached from the cell culture surface using Trypsin (Lonza) and stored at -80 °C until further use.

BJ-EHT vRAS12 cells, treated with tamoxifen, were lysed using 1% sodium deoxycholate lysis buffer as described previously (Post et al., 2017). Proteins were digested overnight in parallel using one of the four following proteases: Trypsin/LysC, GluC, Chymotrypsin or AspN at 1:100 (w/w) at 37 °C. Samples were acidified and desalted using Sep-Pak C18 cartridges (Waters), eluted with 80% acetonitrile (ACN)/0.1% trifluoroacetic acid (TFA) and dried down. Samples were fractionated on a high-pH reversed-phase C18 column (Kinetex 5u Evo C18 100A, 150 × 2.1mm, Phenomenex) coupled to an Agilent 1100 series HPLC over a 60 min gradient. Fractions were concatenated to five samples before phosphopeptide enrichment. Enrichment of phosphorylated peptides was performed on the AssayMap BRAVO platform (Agilent Technologies) using Fe(III)-IMAC cartridges following the method described previously (Post et al., 2017).

Samples were analysed by nanoLC-MS/MS on a Q Exactive HF-X mass spectrometer (Thermo Scientific) equipped with an Agilent 1290 LC system with an LC gradient of 115 min gradient (9% to 35% B). MS settings were as follows: full MS scans (375-1600 m/z) were acquired at 60,000 resolution with an AGC target of 3e6 charges and max injection time of 20 msec. HCD MS2 spectra were generated for the top 12 precursors using 30,000 resolution, 1e5 AGC target, a max injection time of 50 msec, a scan range of 200-2000m/z and normalised collision energy of 27%. MS2 isolation windows were 1.4m/z.

Raw data files were processed with MaxQuant v1.6.3.4 using a Uniprot human database. Carbamidomethyl (C) was set as a fixed modification and Oxidation (M), Acetylation (protein N-terminus) and Phosphorylation (STY) were set as variable modifications. Each proteolytic enzyme was searched individually. Max missed cleavages was set to four for GluC and Trypsin/LysC and 5 for Chymotrypsin and AspN. Results were filtered using a 1% FDR cut off at the PSM level.

### Selection of phosphopeptides for SRM assay

Relevant phosphoproteins and phosphosites within the mTOR pathway were identified by literature search. Phosphopeptides that represented these biologically relevant phosphosites were selected in the following manner: firstly, phosphopeptides were selected from our phosphopeptide library. If multiple phosphopeptides were available for the same phosphosite, the most suitable phosphopeptide was chosen. Firstly, peptides without methionine were favoured over peptides including a methionine. Furthermore, the phosphopeptide with the highest intensity was chosen. If our phosphopeptide library did not yield a phosphopeptide to cover a phosphosite of interest we extracted suitable peptides from publically available databases (Giansanti et al., 2015; Lawrence et al., 2016).

### Generation of spectral library

Selected phosphopeptides were synthesised incorporating an isotopically labelled amino acid. These heavy labelled peptides were used to generate a spectral library. For this, peptides were dissolved and mixed with iRT peptides for retention time alignment (Escher et al., 2012). Peptides were analysed by nanoLC-MS/MS on a Q Exactive HF-X mass spectrometer (Thermo Scientific) as described previously with the following changes: HCD MS2 spectra were generated for the top 15 precursors and max injection time was set to 54 msec. Raw files were subjected to database search using Proteome Discoverer (2.2). Tryptic or GluC were selected as proteolytic enzyme, max 3 missed cleavages were allowed, Carbamidomethyl (C) was set as a fixed modification and Oxidation (M), Acetylation (protein N-terminus) and Phosphorylation (STY) were set as variable modifications. Results were filtered using Percolator, FDR was set to 1%. The resulting files were used to build a spectral library in Skyline.

### SRM assay development

Heavy labelled standard peptides were used to develop the SRM assay. For each precursor, at least three fragments were measured. The topmost intense fragment ions were chosen and complemented with an additional transition if Y1-ion was amongst the top three most intense ions or if an extra transition was needed to distinguish phospho-isomers. Collision energy optimisation was performed for each transition individually as described previously (MacLean et al., 2010).Retention time scheduling was performed using iRT as described previously (Escher et al., 2012).

### SRM assay characteristics

The SRM assay we developed had the following characteristics: LC-MS/MS analysis was performed on an Ultimate 3000 RSLCnano System (ThermoScientific) coupled to a TSQ Altis Triple Quadrupole (ThermoScientific). Peptides were reconstituted in 20mM citric acid, loaded on a pre-column (C18 PepMap100, 5 μm) and separated on a PepMap RSLC C18 column (2μm, 75μm x 25cm) using a 100 min gradient (2.2% to 34% Buffer B 100% ACN + 0.1% FA) at a flow of 300 nl/min. Retention time windows were set to 5 minutes, Q1 and Q3 resolution was set to 0.7 and a cycle time of 5 seconds was used.

### Sensitivity test

Sensitivity of the method was determined by a dilution series of heavy labelled standards spiked into a HELA phosphopeptide background. HELA cells were cultured in DMEM Cells were washed with ice cold PBS, detached from the cell culture surface using Trypsin (Lonza) and stored at -80 °C until further use. Cell lysis, protein digestion (using Trypsin/ LysC) and sample clean-up was performed as described previously for BJ-EHT cells. After sample clean-up, 200μg of cell digest was enriched for phosphorylated peptides (as described previously) and spiked-in with decreasing amount of heavy labelled standards (100 fmol, 50 fmol, 10 fmol, 5 fmol, 1 fmol, 0.1 fmol). This was performed in triplicates. Samples were subjected to the SRM assay described previously. Raw data were analysed using skyline and peaks were individually assessed. Quantification was based on the total peak area.

### MCF10A experiments

MCF10A cells were cultured in DMEMF12 medium, supplemented with 5% horse serum, 1% penicillin/streptomycin, EGF (20ng/ml), hydrocortisone (0.5 mg/ml), cholera toxin (100ng/ml) and insulin (10μg/ml) (normal growth conditions). Cells were then starved from either growth factors (using regular DMEMF12 without additives), amino acids (using DMEMF12 without amino acids), or glucose (using DMEMF12 without glucose) for 24 hours. Afterwards, cells were treated with either growth factors (EGF 20 ng/ml and insulin 10 μg/ml), amino acids or glucose for different time points. Cells were washed with ice cold PBS, detached from the cell culture surface using Trypsin (Lonza) and stored at -80 °C until further use. Cells were lysed as described previously for BJ-EHT cells. Protein digestion was performed using Trypsin/LysC or GluC (1:75 w/w enzyme: substrate ratio) overnight at 37 °C. Samples were acidified and tryptic and GluC peptides were combined. Samples were desalted using the AssayMAP Bravo Platform (Agilent Technologies) using C18 cartridges (Agilent Technologies). Cartridges were primed using 100 μl priming solution (80% ACN 0.1% FA) and subsequently equilibrated using 50 μl equilibration solution (0.1% FA in miliQ). Samples were loaded at 10 μl/min and eluted with 50 μl of priming solution at 10 μl/min. After desalting, heavy labelled standards were spiked into each sample. Subsequently, enrichment of phosphorylated peptides was performed on the AssayMap BRAVO platform (Agilent Technologies) using Fe(III)-IMAC cartridges following the method described previously (Post et al., 2017). Samples were dried down using and stored at -80°C until they were subjected to LC-MS analysis (as described previously under SRM assay characteristics).

