## Supplementary Figures for "Capturing the signalling dynamics of the MAPK-AKT-mTOR pathway in a single targeted phosphoproteomics assay"

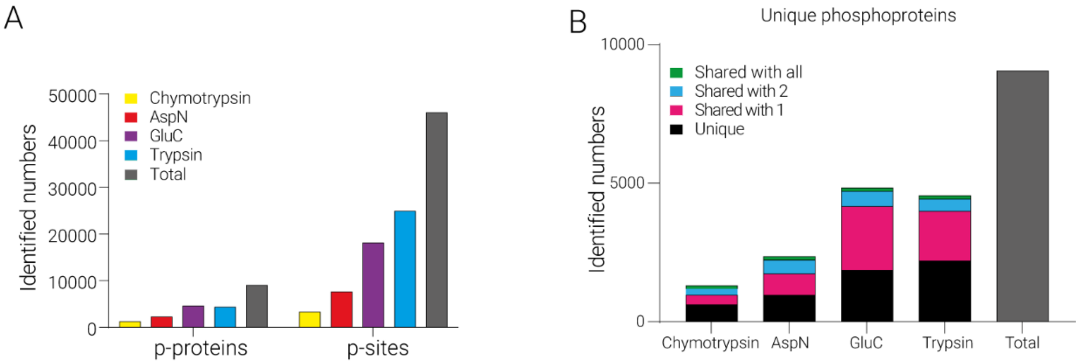


**Supplementary Figure 1. In-house built phosphopeptide library numbers.** (A) Number of identified phosphoproteins, phosphopeptides and phosphosites per protease. (B) Phosphoproteins identified per protease coloured by uniqueness.

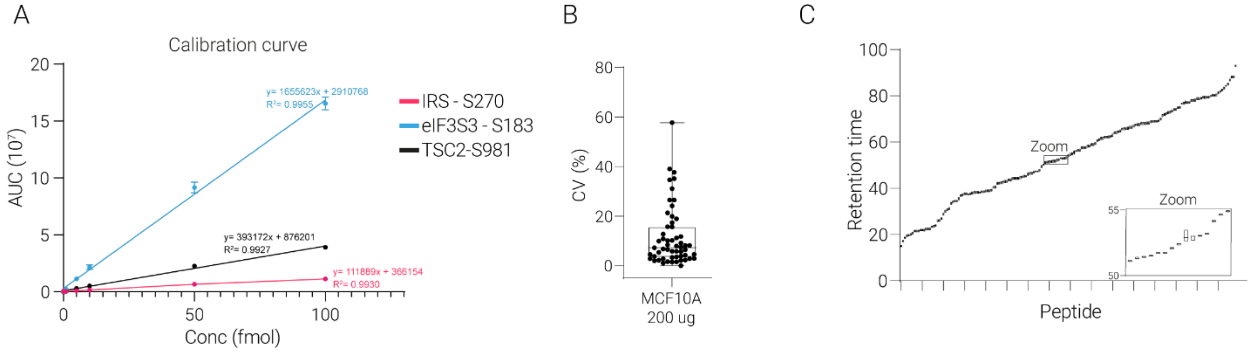


**Supplementary Figure 2. Analytical characteristics of the targeted MS assay.** (A) Calibration curve of three phosphopeptides titrated down in a phosphopeptide background illustrates large dynamic range. (B) Coefficient of variation for replicate measurements illustrates quantitative reproducibility. (C) Retention time reproducibility is high, demonstrating robust LC setup.

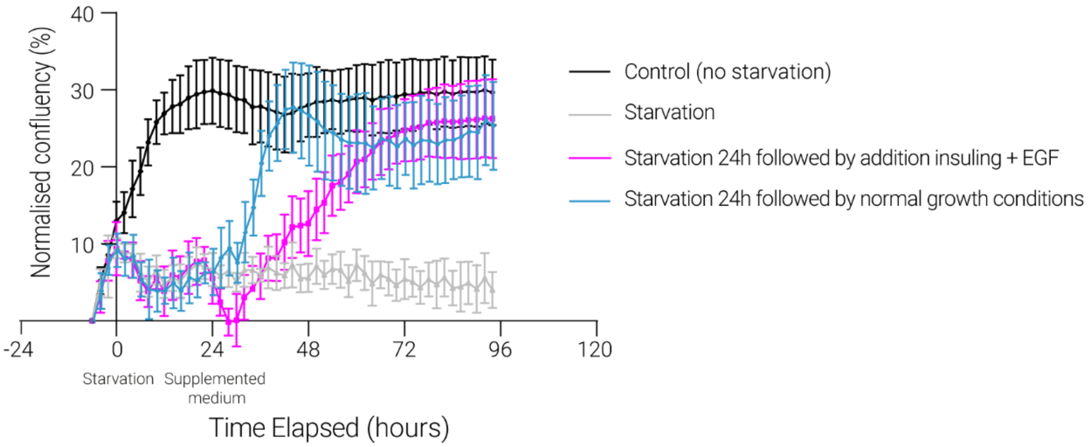


**Supplementary Figure 3. Cellular growth curves.** Cellular growth curves under normal growth conditions (black), continues starvation (grey), 24h starvation followed by addition of EGF and insulin (pink) and 24h starvation followed by normal growth conditions (blue).
